# Computationally Guided Design of Novel Selective Serotonin Reuptake Inhibitors

**DOI:** 10.1101/2020.08.21.262006

**Authors:** Kavya G. Achyutuni, Sambid Adhikari, Peishan Huang, Justin B. Siegel

## Abstract

Current selective serotonin reuptake inhibitor (SSRI) drugs such as paroxetine, fluoxetine and escitalopram are prescribed for mental health conditions such as anxiety and depression. However, there are still several unpleasant side effects which fuels the need for new designs. In this work, computational studies were conducted to design two novel drug candidates with improved docking scores than paroxetine, an existing SSRI drug. Homology analysis was carried out to determine the suitability of an organism for preclinical trials and revealed laboratory mouse (*Mus musculus*) to be a good candidate.

## INTRODUCTION

Depression is a mental health disorder that affects around 16 million adults in the United States every year and is the third most common cause for hospitalization.^1^ The increase in prevalence of depression has led to the development of several drugs by pharmaceutical industries. Currently, the most common way to target depression is by inhibiting the binding of serotonin to the ts3 human sodium-dependent serotonin transporter.^2^ Serotonin is a neurotransmitter responsible for wellbeing and happiness and is the natural substrate of the serotonin transporter.^2^ The serotonin transporter facilitates the reuptake of extracellular serotonin, thereby terminating neurotransmission.^2^ The lack of extracellular serotonin is thought to cause neuropsychiatric disorders such as anxiety and depression.^2,3^ Thus, selective serotonin reuptake inhibitors (SSRIs) are a class of drugs that have been developed to compete with the binding of serotonin. The binding of SSRIs to the transporter in place of serotonin increases the levels of extracellular serotonin and is thought to improve mood.^2^ The serotonin transporter is therefore a good target for antidepressants.

Several SSRI drugs such as citalopram, escitalopram, fluoxetine, fluvoxamine and paroxetine have been successful in treating anxiety and depression.^4^ These drugs bind to the serotonin transporter thereby preventing serotonin from binding.^2^ Although these drugs have been effective, there are certain pharmacokinetic and pharmacodynamic properties that have negative effects on the body. For example, drugs such as fluoxetine and paroxetine contain several toxicophores which are functional groups that can transform into unstable intermediates in the body and lead to abnormal cellular function.^5^ The half-life of drugs like fluoxetine have been found to be three days, which can be toxic to the body.^6^ Additionally, some of the common side effects of the drugs listed include headache, nausea, and constipation.^7^ Drugs such as paroxetine have been shown to have negative effects on the elderly and patients with renal and hepatic disease.^8^ As a result, new and improved designs are needed to potentially minimize these side effects.

The starting point for new designs begins with the identification of the pharmacophore (Fig. 1A), a set of functional groups common to all SSRIs.^9^ Although novel drugs may contain different functional groups compared to existing drugs, the pharmacophore is essential for activity.^9^ A previous study proposed a pharmacophore model that contains two aromatic rings, one hydrophobic group and one positive ionizable group (Fig. 1A).^9^ The study also stated that cation-pi interactions could play a role in stabilizing the molecule in the active site by binding to nearby residues (Fig. 1B).^9^ Using this pharmacophore model as a starting point, two novel SSRI drug candidates were proposed using computational and chemical intuition-guided methods. These two novel drug candidates were compared to existing SSRI drugs based on ADMET (absorption, distribution, metabolism, excretion, toxicity) properties, and docking scores. Both drug candidates showed significantly improved docking scores, better predicted oral bioavailability and fewer toxicophore groups compared to paroxetine, the model for the study. Additionally, homology analysis was performed to select an organism for clinical trials to study how the drug candidates can potentially interact with the body.

**Figure 1.**
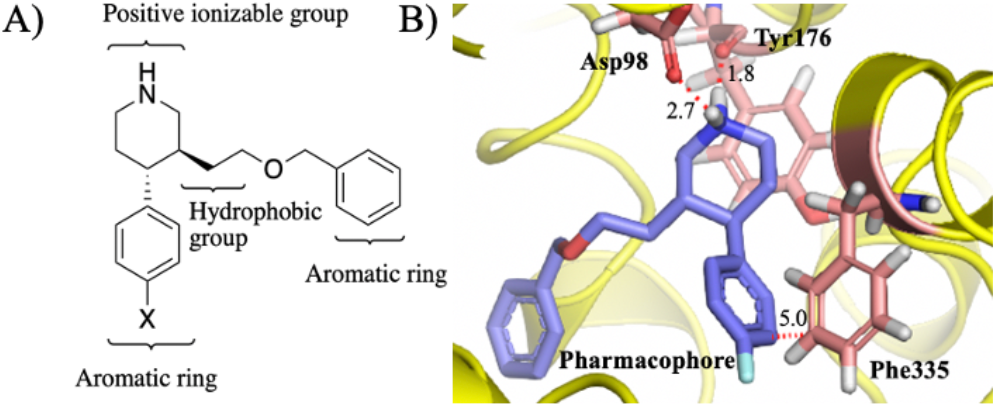
A) Proposed pharmacophore model of SSRI showing a positive ionizable, aromatic, and hydrophobic groups. B) Molecular interactions (red dotted lines) between pharmacophore (blue) and Asp98, Tyr176, Phe335 residues of ts3 serotonin transporter.

## METHODS

The crystal structure of the ts3 human serotonin transporter with reversible inhibitors bound to the active site (PDB ID: 5I71)^12^ obtained from Protein Data Bank was used for drug design, molecular simulation, and docking. Using the OpenEye software suite, MakeReceptor^20^ was used to define the binding site on the serotonin transporter. To design a drug candidate based on the pharmacophore using bio-isosteric method, vBROOD^17^ was used to generate replacements of functional groups from the model, paroxetine. Additionally, SciFinder was used to confirm that the molecules were not previously studied. Once the ideal candidates were designed, Gaussview^23^ and Gaussian 09^14^ were used to build and optimize drug candidates, respectively. Conformer libraries of known drug molecules and designed candidates were generated using OpenEye OMEGA^22^, after which the conformer libraries were docked into the active site using FRED. ^21^

ADMET properties such as hydrogen bond donors and acceptors, molecular weight, number of chiral centers and xLogP bioavailability values were found using OpenEye FILTER.^16^ PyMOL^15^ was used to visualize the protein-ligand interactions as well as finding the measurements between the ligand and residues. BLAST^18^ was used to search for human serotonin transporter protein homologs. Finally, Jalview^19^ was used to analyze sequences of the serotonin transporter of various organisms for homology analysis.

## RESULTS AND DISCUSSION

### Evaluation of known SSRI drugs

Four known SSRI drugs shown in Table 1 were evaluated based on their molecular interactions, ADMET properties, and docking scores. Paroxetine was found to have the most negative docking score as well as two aromatic rings and a positive ionizable group. Additionally, based on the ADMET properties generated by OpenEye FILTER (Table 1), all four drugs followed Lipinski’s rule of five.^24^ All four molecules were also found to have two or less chiral centers. The xLogP value of paroxetine was 3.37, which indicates that paroxetine can be administered orally. The other three drugs were found to have xLogP values greater than four, indicating that they need to be administered transdermally. Transdermal drug administration has been found to have variability in the amount of drug absorbed into the bloodstream based on biological factors such as sex, race and age. ^10^ Hence, in this work, the oral method of administration is preferred.

**Table 1.** Docking scores and ADMET properties of known SSRI drugs Paroxetine, Fluoxetine, Fluvoxamine, and Escitalopram.

| Name | Docking Score | Molecular weight (g/mol) | Lipinski H-bond acceptors | Lipinski H-bond donors | xLogP | # Chiral centers |
| --- | --- | --- | --- | --- | --- | --- |
| Paroxetine | -14.87 | 330.37 | 4 | 2 | 3.37 | 2 |
| Fluoxetine | -13.91 | 310.33 | 2 | 2 | 4.66 | 1 |
| Fluvoxamine | -13.88 | 319.34 | 4 | 3 | 4.07 | 0 |
| Escitalopram | -13.69 | 325.40 | 3 | 1 | 4.18 | 1 |

Although the four drugs have been administered successfully, there have been side effects in patients due to the presence of toxicophore groups. For example, in paroxetine (Fig. 2A), the fluorophenyl and benzodioxol groups are known toxicophores and can cause abnormal cellular function.^5^ Despite the presence of toxicophore groups, paroxetine has the lowest docking score of −14.87 and the best oral bioavailability. Since improvement on this drug can further lower the overall docking scores compared to the other three drugs, paroxetine was chosen to be the starting point for the design of novel SSRI drugs.

**Figure 2.**
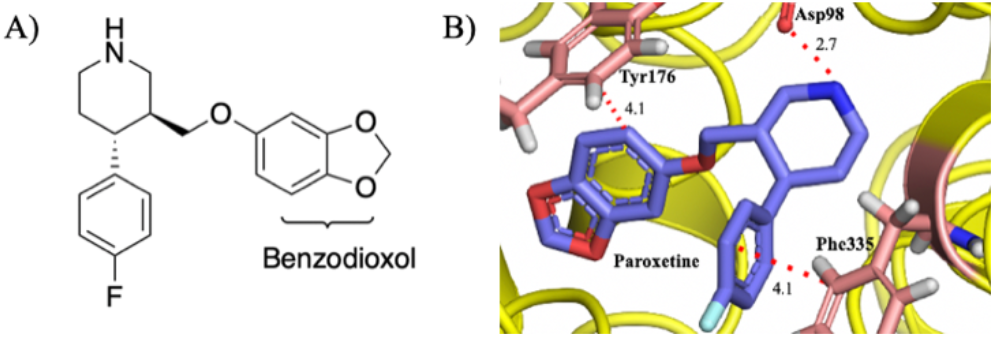
A) 2D image of Paroxetine. B) Interactions (red dotted lines) between Paroxetine (blue) and Asp98, Tyr176, Phe335 residues of ts3 serotonin transporter.

Using PyMOL as a visualization tool, several key interactions are predicted to be important. Paroxetine is non-covalently bound to the active site of the serotonin transporter shown in Figure 2B. Some of the strongest interactions include two pi-pi stacking interactions between the fluorophenyl group and the aromatic ring of the Phe335 side chain at 4.1 Å, and between Tyr 176 and the benzene ring of the benzodioxol group at 4.1 Å. A strong hydrogen bond between the hydrogen of the piperidine and the carbonyl of the Asp98 side chain at 2.7 Å was also observed. Based on the pharmacophore model, docking scores and interactions with active site residues, two new drug candidates were designed.

### Drug Candidate 1 designed using computational studies

The first drug candidate (Fig. 3A) was proposed using computational studies that began by loading paroxetine into vBROOD. Several candidates which contained bio-isosteres as replacements for the toxicophoric benzodioxol group were generated. The conformer library of those candidates was generated and it was subsequently docked into the human receptor. The molecule with the best docking score of −18.65 was named Candidate 1, which is a 25.42% better score compared to paroxetine (Table 1). This new molecule contained a carbonyl group in place of the ether, and a pyrimidine ring attached to a five-membered ring in place of the benzodioxol group.

**Figure 3.**
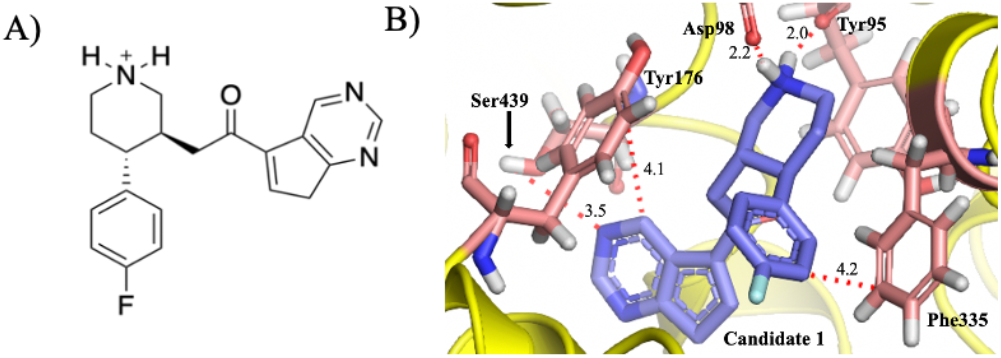
A) 2D image of proposed Candidate 1 using bio-isosteric method with pyrimidine ring attached to five-membered ring. B) Interactions (red dotted lines) between Candidate 1 (blue) and Asp98, Tyr176, Phe335, Tyr95, Ser439 residues of ts3 serotonin transporter.

While most of the interactions remained similar as with paroxetine, Candidate 1 has a more negative docking score due to a new hydrogen bond interaction at 3.5 Å between the nitrogen of the pyrimidine ring and the hydrogen of Ser439. Additionally, the hydrogen bond distance between Asp98 and piperidine, became 0.5 Å shorter compared to paroxetine, thereby making the bond stronger. The new interaction and shorter hydrogen bond contribute to the overall more negative docking score of Candidate 1.

To determine the how the drug could potentially interact with the body, ADMET screening was carried out. The screening revealed that the molecular weight of Candidate 1 was below 500 g/mol and there were an acceptable number of Lipinski hydrogen bond donors and acceptors (Table 2). Although the xLogP value decreased compared to paroxetine, Candidate 1 is still in the orally administrable range which is around 0-3. Additionally, two chiral centers were present and the pyrimidine ring attached to the five-membered ring is not known to be a toxicophore, unlike the benzodioxol group of paroxetine. Based on the elimination of the toxicophore and the improved docking score, Candidate 1 is expected to be a promising drug candidate.

**Table 2.** ADMET properties of Candidate 1 and Candidate 2.

| Name | Molecular Weight (g/mol) | Lipinski H-bond acceptors | Lipinski H-bond donors | xLogP | # Chiral centers |
| --- | --- | --- | --- | --- | --- |
| Candidate 1 | 338.40 | 4 | 2 | 1.78 | 2 |
| Candidate 2 | 354.40 | 5 | 3 | 1.28 | 2 |

### Drug Candidate 2 designed using chemical intuition

The design of drug candidate 2 was based off of Candidate 1. Since Candidate 1 had a weak hydrogen bond between Ser439 and the aromatic ring at 3.5 Å, the intuition behind designing drug candidate 2 was the possibility of another stronger hydrogen bond to increase binding to the active site. A hydroxyl group was added to the second carbon from the carbonyl group onto the five-membered ring (Fig. 4A). This molecule was built in Gaussview and optimized in Gaussian. The conformer library of this novel compound was docked into the human receptor. This molecule, named Candidate 2, was found to have a docking score of −17.41 which is a 17.08% improvement compared to paroxetine.

**Figure 4.**
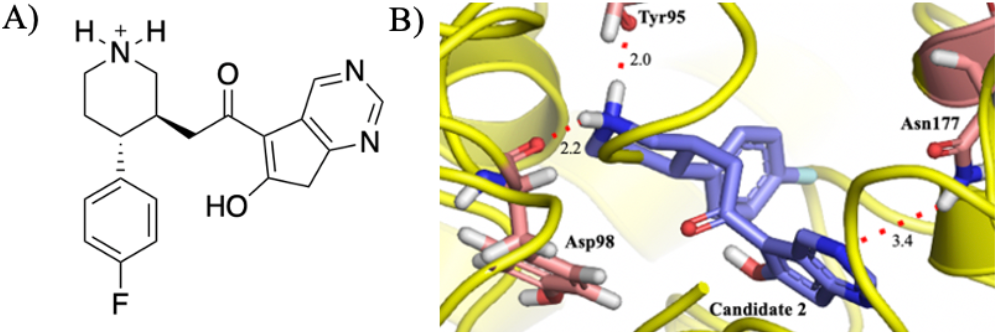
A) 2D image of proposed Candidate 2 using chemical intuition method with addition of hydroxyl group. B) Interactions (red dotted lines) between Candidate 2 (blue) and Asp98, Tyr95, Asn177 residues of ts3 serotonin transporter.

To further understand why Candidate 2 has a better docking score, its interactions with the ts3 serotonin transporter were shown in Figure 4B. Although most interactions remain the same as in Candidate 1, the weak hydrogen bond between the nitrogen of the pyrimidine ring and the hydrogen of Ser439 at 3.6 Å has been replaced by a stronger hydrogen bond between nitrogen of the six-membered ring and Asn177 at 3.4 Å. This novel hydrogen bond is facilitated by the addition of the hydroxyl group on the molecule, which pushes the aromatic ring system further outward, closer to the residue. The pi-pi stacking interactions between Phe335 and the fluorophenyl group and between Tyr176 and the pyrimidine ring, however, have been lost. This loss is possibly due to a steric clash that is causing the OH group to move away from the fluorophenyl group thereby causing the conformation of the aromatic rings to rotate, and explains why Candidate 2 has a better docking score than paroxetine but worse than that of Candidate 1.

Finally, the ADMET properties of Candidate 2 were evaluated (Table 2). The molecular weight of this molecule was less than 500 g/mol, and there were an acceptable number of Lipinski hydrogen bond donors and acceptors. As expected, the number of hydrogen bond donors and acceptors both increased by one due to the addition of the alcohol group. The xLogP value is much lower than that of paroxetine and also slightly lower than that of Candidate 1. However, it remains a good candidate for oral administration. Additionally, two chiral centers are present and the five and six-membered ring systems along with the hydroxyl group are not known to be toxicophores. Similar to Candidate 1, this improvement also eliminates the toxicophore from paroxetine. The ADMET properties and docking score confirm that this molecule should be further investigated as a potential lead compound.

On the whole, Candidate 1 and Candidate 2 showed improved docking scores and hence improved binding to the active site of the serotonin transporter. This is due to novel hydrogen bond interactions between the active site and both candidates. Since the benzodioxol group has been replaced, both candidates have one less toxicophore than paroxetine. However, the fluorophenyl group on both candidates still remains a toxicophore. Both drug candidates have just two chiral centers and the xLogP values of both drugs indicate that they can be administered orally. This confirms that both drugs would make potential good candidates for further study.

### Selection of an organism for preclinical trials based on homology analysis

In order to examine the viability of these drug candidates to become potential drugs available in the market for treating depression, preclinical trials are an important step. In order to perform preclinical trials, an organism with a protein homologous to the human target should be present in order to evaluate toxicity as well as potential efficacy. Based on a BLAST search, the results indicated that the sodium-dependent serotonin transporter of *Mus musculus* commonly known as laboratory mouse is homologous to the human sodium-dependent serotonin transporter with a 99% query cover and 100% identity.

The sequence of the human ts3 serotonin transporter was compared to the homolog sequences. It was observed that the residues Tyr95 and Asp98, which interact with both Candidate 1 and Candidate 2, were conserved in the mouse serotonin transporter with 100% identity. Additionally, Phe335 and Tyr176 residues which interact specifically with Candidate 1 and Asn177 which interacts with Candidate 2 were also conserved with 100% identity. Active site residue Ser439, however, was not conserved in the mouse serotonin transporter and a threonine residue was observed instead. Although Ser439 is replaced by Thr439, both residues have an alcohol group which can still provide potential hydrogen bond interactions. To understand how the candidates interact with the murine ts3 receptor compared to the human ts3 receptor, the candidates were docked into the murine transporter (PDB ID: 6DZW)^11^ and their interactions were shown figure 5. Most distances between the interacting groups were similar to the human transporter. The OH group of Thr439 not only interacts with Candidate 1 but also forms a stronger hydrogen bond with the nitrogen of the six membered ring at 3.4 Å (Fig. 5), compared to that between the drug and the Ser439 of the human serotonin transporter at 3.6 Å. Based on similar residues shown between human and mouse serotonin transporters, this organism should be viable model for preclinical trials.

**Figure 5.**
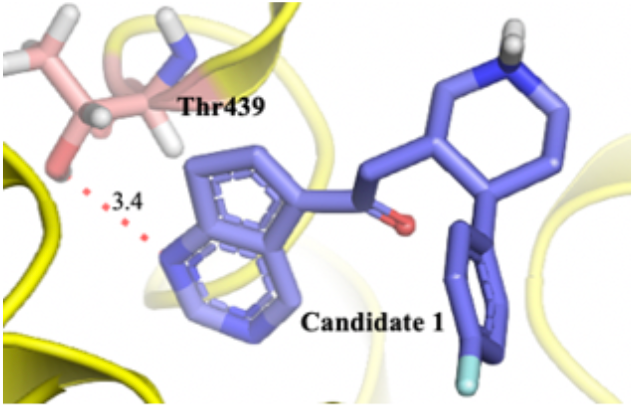
Interactions (red dotted lines) between Candidate 1 (blue) and Thr439 residue in *Mus musculus* serotonin transporter.

## CONCLUSION

Current serotonin reuptake inhibitors have some undesirable side effects such headache, nausea and constipation as well as the presence of toxicophore groups which can cause abnormal cellular function. Thus, there is a need for new drug candidates that can mediate these problems. In this work, two drug candidates were proposed and had significantly better docking scores compared to paroxetine while still meeting ADMET criteria. Further homology analysis using the *Mus musculus* serotonin transporter as the homolog reveals similar active-site residues that are found to be important for binding with drug candidates, thus, provides a suitable model for preclinical trials.

Although these computational tools proved useful for finding potential candidates in a quick and efficient manner, and the results are promising in the development of new drugs to combat depression and anxiety, preclinical and clinical trials need to be conducted in order to determine whether these molecules provide the desired outcomes in a biological environment. Despite producing good docking scores and suitable ADMET properties, these drugs could still react with more than one target in the body and produce side effects. The next step, therefore, is to test these two compounds to determine toxicity and subsequently efficacy profiles in animal models. ^25^

## Author Contributions

Research was designed by all authors; all experiments were carried out by L.C.R. The manuscript was written through contributions of all authors. All authors have given approval to the final version of the manuscript.

## ACKNOWLEDGMENT

Research reported in this publication was supported by UC Davis, the National Science Foundation Award Numbers 1827246, 1805510, 1627539, the National Institute of Environmental Health Sciences of the National Institutes of Health (NIH) under Award Number P42ES004699, UC Davis, NIH Award Number R01 GM 076324-11 and the Rosetta Commons. The content is solely the responsibility of the authors and does not necessarily represent the views of the National Institutes of Health or National Science Foundation. This study was derived from a course based undergraduate research study conducted in Chemistry 130B at UC Davis.

